# Rapid exchange of stably bound protein and DNA cargo on a DNA origami receptor

**DOI:** 10.1101/2022.01.16.476526

**Authors:** James W.P. Brown, Rokiah G. Alford, James C. Walsh, Richard E. Spinney, Stephanie Y. Xu, Sophie Hertel, Jonathan F. Berengut, Lisanne M. Spenkelink, Antoine M. van Oijen, Till Böcking, Richard G. Morris, Lawrence K. Lee

**Author notes:** These authors contributed equally (James W.P. Brown, Rokiah G. Alford).

## Abstract

Biomolecular complexes can form stable assemblies yet can also rapidly exchange their subunits to adapt to environmental changes. Simultaneously allowing for both stability and rapid exchange expands the functional capacity of biomolecular machines and enables continuous function while navigating a complex molecular world. Inspired by biology, we design and synthesize a DNA origami receptor that exploits multi-valent interactions to form stable complexes that are simultaneously capable of rapid subunit exchange. The system utilizes a mechanism first outlined in the context of the DNA replisome, known as multi-site competitive exchange, and achieves a large separation of time scales between spontaneous subunit dissociation, which requires days, and rapid subunit exchange, which occurs in minutes. In addition, we use the DNA origami receptor to demonstrate stable interactions with rapid exchange of both DNA and protein subunits, thus highlighting the applicability of our approach to arbitrary molecular cargo; an important distinction with canonical toehold exchange between single-stranded DNA. We expect this study to be the first of many that use DNA origami structures to exploit multi-valent interactions for the design and synthesis of a wide range of possible kinetic behaviors.

## INTRODUCTION

Multi-protein complexes are essential for performing many tasks that are fundamental to living organisms, including but not limited to the replication^1^ and transcription^2^ of DNA, respiration^3^, cell division^4^, cell motility^5^ and signaling^6^. These complexes can be highly stable and robust, continuously functioning over extended timescales on the order of hours and with great precision. Additional layers of regulation allow subunits to be rapidly exchanged in response to changes in the intracellular or extracellular environment^7^. An example is the bacterial flagellar motor, which functions as a stable multi-protein complex that rotates flagellar filaments to propel certain bacteria through viscous media. Yet subunits in the flagellar motor rapidly exchange with those in solution in response to changing chemical gradients^8^, to switch directions^9, 10^ or to remodel and tune its detection limits^11^ or power output^12, 13^. Another example is the replisome, the multi-protein complex responsible for DNA replication, which readily changes stability depending on the availability of components in its environment^14–19^.

Subunits in these multi-protein complexes thus have apparently competing requirements: they must bind with sufficient stability to robustly perform their function but then allow for rapid unbinding for subunit exchange in response to changes in the environment. This apparent paradox, known as stability vs exchange^20^ can be illustrated in the following simple reaction where a receptor molecule R, is in a stable heterodimeric complex with an incumbent cargo molecule I, which is then replaced by a freely diffusing competitor cargo molecule C, to form the complex RC:

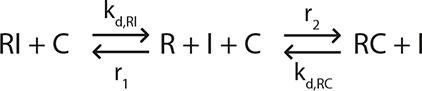

Stability of the RI complex is inversely proportional to the dissociation rate constant of the RI complex (*k_d,RI_*). However, where the RI interaction is a simple monovalent interaction, the incumbent molecule must first dissociate from the receptor before the competitor can bind to form the RC complex. Thus, a slow *k_d,RI_* dissociation rate constant required for the stability of RI, precludes the rapid exchange of an incumbent molecule with a competitor molecule, irrespective of its association rate with the receptor, which is determined by the association rate constant (*k_a,RC_*) multiplied by the concentrations of R and C (*r*_2_ = *k_a,RC_*[*R*][*C*]). Hence the requirements of stability and rapid exchange act in direct competition.

Subunits however, need not bind via monovalent interactions. In biology, multi-protein complexes are often formed through a number of weak multivalent interactions which, in a process known as avidity, allows for high stability ^21–24^. To illustrate this, we can then consider a bivalent system in which a receptor molecule contains a distinct primary (1) and secondary (2) binding site, which bind to complementary distinct 1* and 2* binding sites on a cargo molecule respectively (Figure 1A).

**Figure 1.**
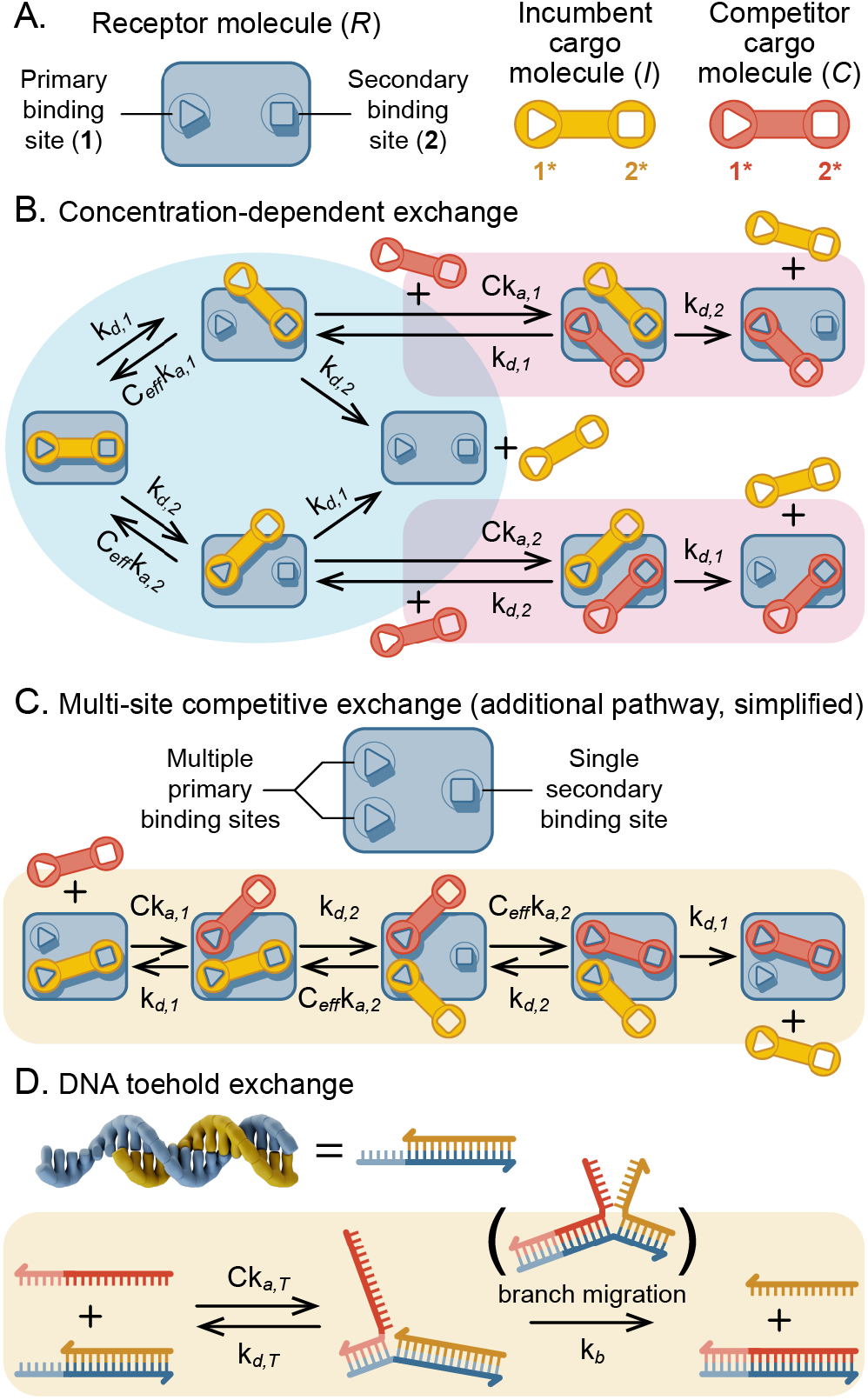
Reaction pathways for concentration-dependent exchange of molecular cargo. (A) Depiction of a system that forms heterodimers via a bivalent interaction with independent primary and secondary binding sites. (B) Pathways for spontaneous dissociation (blue shading) and concentration-dependent exchange (pink shading) for a standard bivalent interaction. (C) Additional exchange pathways in the presence of an additional primary binding site to enable multi-site competitive exchange. (D) Illustration of toehold exchange mechanism for DNA strand displacement where the toehold domain (T, blue) is bound by an invader molecule and the incumbent cargo (bound via a hybridization domain, red) is displaced via branch migration.

Figure 1B depicts an exchange reaction which illustrates (blue shading) how spontaneous dissociation requires both the primary and secondary binding sites to simultaneously unbind for a fully bound cargo to completely dissociate. Moreover, we see that upon unbinding of any single interaction, the free binding sites remain within close proximity, which results in a high local or effective concentration (*C_eff_*). *C_eff_* can be defined mathematically as the ratio of the product of the equilibrium dissociation constants (*K_D_*) of each binding site and the overall equilibrium dissociation constant of the interaction^25, 26^, which for the above reaction equates to:

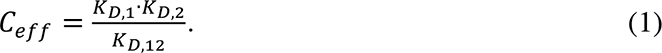

*K_D,1_* and *K_D,2_* are the equilibrium dissociation constants for the primary and secondary binding sites respectively and *K_D,12_* is the equilibrium dissociation constant for the interaction between receptor and cargo molecules, which for a divalent interaction incorporates both binding sites. Thus when *C_eff_* is large, as is typical for intramolecular interactions given their short length scales, bivalency allows for stability to be achieved with relatively weak interactions that may be unstable in isolation. Bivalency also enables concentration-dependent competitive exchange, which has been observed experimentally for the binding of proteins to DNA^27–29^. Competitive exchange occurs because a competitor molecule, which also binds to the primary and secondary sites, can compete with a partially bound cargo molecule for the available binding site without having to wait for complete dissociation (Figure 1B, pink shading). The probability for a competitor molecule to outcompete rebinding of the partially bound incumbent molecule depends on the concentration of free competitor molecules [*C*], thus resulting in a concentration-dependent effect on the dissociation of the initially bound molecule. However, significant competitive exchange therefore also requires that [*C*] approaches or exceeds the typically high *C_eff_*, which can be practically difficult to attain. At lower concentrations of competitor molecule, the rate of dissociation of the incumbent molecule is insensitive to [*C*], and the exchange rate of the system is non-competitive, approximating that of a monovalent interaction (Spinney *et al.* 2022).

An alternative mechanism known as multi-site competitive exchange (MSCE) has recently been proposed^20^ to occur in the DNA replisome in which the incumbent DNA polymerase binds simultaneously to the elongating DNA strand, akin to a single secondary site, and to one of multiple possible primary binding sites on the DNA helicase enzyme^14, 30, 31^. The MSCE reaction mechanism is illustrated in Figure 1C, with a simple receptor configuration consisting of a single secondary site and two primary binding sites. Under dilution, the incumbent cargo molecule binds stably, similar to a regular bivalent interaction. In the presence of free competitor molecules however, unbound primary binding sites can be occupied without competing with the incumbent cargo molecule. The unbound site on the immobilized competitor cargo molecule also has a high local concentration at its unbound receptor site, which assuming equal distance between the secondary site and both primary binding sites, is equal to *C_eff_*. Upon spontaneous dissociation of the incumbent cargo from the secondary site, the immobilized competitor molecule has equal probability of occupying the unbound secondary site, thus achieving a stable multivalent interaction and consequently, spontaneous dissociation of the incumbent cargo is favorable. Importantly, the range of [*C*] through which exchange is sensitive depends on *K_D,1_*, since this determines the probability that the free primary binding site is occupied by a competitor molecule. Thus, when *K_D,1_* ≪ *C_eff_*, concentration-dependent exchange occurs at bulk concentrations much lower than required for regular subunit exchange.

DNA hybridization is another example of a well-defined multivalent interaction in which two complementary strands form a single duplex via multiple, weak base-pairing interactions. This multivalency has been widely exploited to achieve stable interactions while allowing for rapid DNA strand displacement in a mechanism known as toehold exchange^32, 33^. The toehold exchange mechanism is illustrated in Figure 1D and involves a DNA duplex with one longer strand and one shorter strand, which in the notation above, are equivalent to the receptor and incumbent cargo molecule respectively. The additional unpaired DNA bases in the receptor are referred to as the toehold and act as an accessible binding site for free competitor cargo DNA strands. Thus, as with the primary binding sites in MSCE, toehold sites are occupied in a concentration-dependent manner that is most sensitive to the equilibrium dissociation constant of the toehold (*K_D,T_*). Once bound to the toehold, there is a significant probability that the competitor strand will consecutively displace base pairing interactions in the incumbent strand in a process termed branch migration. Branch migration has a characteristic rate constant (*k_b_*), which is dependent on the length of the DNA strand that is hybridized to the incumbent strand. The kinetics of toehold exchange have also been explored in detail^34–37^. DNA strand exchange rates and the stability of the toehold domain, which varies approximately exponentially with the binding strength^37, 38^, are tightly coupled. Moreover, exchange rates saturate at toehold lengths of 6-10 nucleotides^37, 38^. Remote toehold exchange enables greater control over DNA strand displacement rates by introducing a single-stranded DNA spacer between the toehold and displacement domain^38^. The spacer serves to decouple the toehold and displacement domains and hence, as with MSCE, results in two distinct binding sites, whose proximity can be described with *C_eff_*. *C_eff_* can be controlled by altering the properties of the spacer to alter branch migration rates with minimal effect on toehold association rates ^38^.

Toehold exchange has been used extensively to actuate the motions of dynamic DNA nanostructures^32, 33, 39–49^. However, predictive models of toehold exchange rely on empirical terms^27^ for *k_b_* or, in the case of remote toehold exchange, blunt-ended strand exchange^42^. Consequently, the kinetics of rationally designed systems are difficult to predict. Toehold exchange is also limited to the exchange of hybridized DNA strands, whereas DNA nanostructures are increasingly being utilized to control the spatial localization of nanoparticles^50^ or other biomolecules such as proteins^51^. In contrast to toehold exchange, MSCE offers a more general and predictable mechanism for stability vs exchange that is applicable to a wide variety of reversible chemistries. Here, we construct the first engineered MSCE system for the exchange of both DNA and protein molecules on a DNA origami receptor. We show that the kinetics of such a system can be entirely predicted by a model of MSCE and that the DNA origami receptor can act as a general receptor for multiple substrates of interest. This study thus acts as a proof-of-principle for the MSCE model, which to our knowledge has not been directly tested in an experimental system. The use of DNA nanostructures for harnessing multivalent interactions can readily be generalized to enact a vast array of interesting rationally designed kinetic properties, which we explore theoretically in a separate publication (Spinney *et al.* 2022). We therefore expect this study to be of broad interest within the field of biomolecular and nanoscale design where the controllable exchange of functional molecules is of significant interest.

## RESULTS

### Design and synthesis of synthetic multi-site competitive exchange system

In order to develop a system capable of the concentration dependent exchange of molecular cargo, a 3D DNA origami structure^52, 53^ was designed and built to act as a receptor containing multiple binding sites for a DNA or protein cargo of interest. The DNA origami structure was designed to fold into a rectangular prism (L × W × H = 49.6 × 25.8 × 18.3 nm) comprising a 12 × 5 honeycomb lattice of DNA helices running along its length. A portion of the middle of the top 3 rows of helices were excluded from the structure, resulting in an accessible cavity with a length of 14.6 nm within which the cargo could bind (Figure 2A). To maximize accessibility, binding sites were located on ends of DNA helices at the outer edges of the cavity (Figure 2B). Binding specificity could be encoded both chemically and spatially. Chemical specificity at each binding site was achieved through the utilization of orthogonal DNA binding sequences through which the binding affinity could also be tuned. Spatial specificity is required to achieve MSCE: the cargo is required to bind to the DNA origami receptor simultaneously via two distinct sites. In keeping with prior nomenclature^20^ we refer to these as the primary and secondary sites (Figure 2B and C). There must also be at least one additional binding site which is distinct but with identical properties to either site. The DNA origami receptor was therefore configured to contain a single secondary binding site in the middle of one side of the cavity, and up to two primary binding sites on the other side of the cavity (Figure 2B). Primary sites were designed to be evenly distributed and equidistant from the secondary binding site. For binding to a DNA cargo, binding sites were 10-nucleotide, single-stranded DNA extensions, the sequence of which initially consisted of 5 × GA repeats and 5 × CA repeats respectively at the primary and secondary sites respectively. This ensured orthogonality and based on our previous SPR measurements^54^, predicted high stability but rapid maximal exchange rates (discussed below). The DNA cargo consisted of a single, rigid DNA duplex with single-stranded DNA extensions on either end that were complementary to the primary and secondary binding sites respectively (Figure 2C). The distances between each primary binding site and the secondary binding site were designed to match the dimensions of the cargo molecule, allowing the cargo to simultaneously access one primary and one secondary site (Figure 2D). In contrast, the distance between primary binding sites were designed to be too close to be simultaneously accessible by both ends of a single DNA cargo molecule.

**Figure 2.**
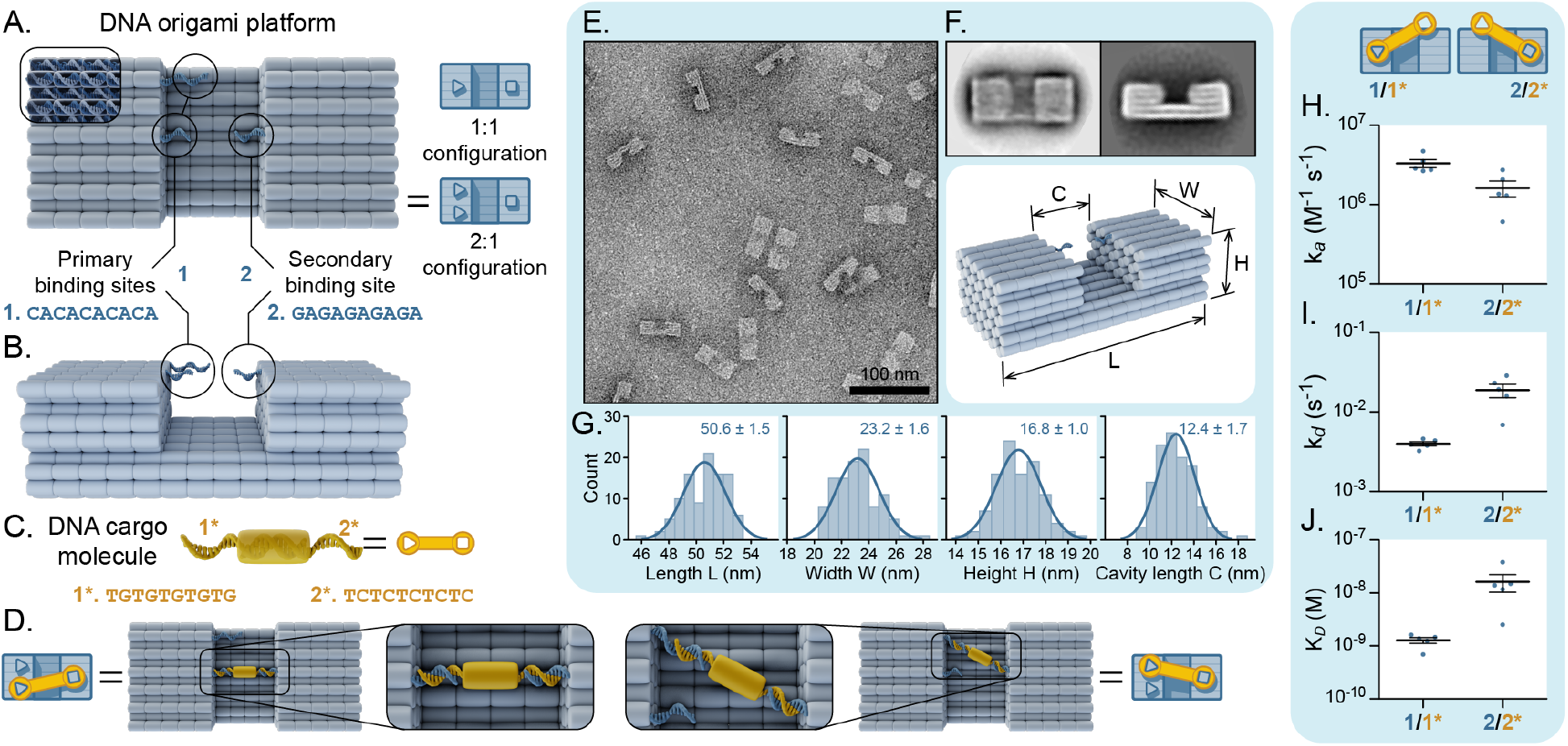
Design and synthesis of a DNA origami based system for multi-site competitive exchange. (A) 3D rendering of the design of the DNA origami receptor with DNA duplexes depicted as cylinders. The DNA origami receptor comprises a rectangular prism with a cavity within which cargo molecules can bind. (B) Location of primary and secondary binding sites on either side of the origami cavity. (C) Design of DNA cargo molecules, consisting of a single hybridized DNA duplex with single-stranded DNA extensions complementary to the primary and secondary binding sites. Staple extensions extend from the DNA origami receptor at the primary and secondary binding sites in a 3’−> 5’ and 5’−> 3’ direction respectively. All sequences are written in a 5’−> 3’ direction. (D) Rendering of DNA origami receptor fully bound to a fluorescent incumbent DNA cargo molecule. (E) Representative wide-field TEM micrograph of purified DNA origami receptor molecules with 2D class averages computed using RELION^55^ in two different orientations shown in (F) where Length, Height, Cavity length and Width are defined (L,H,C and W respectively). (G) Distributions of dimensions taken from >100 individual particles fit with a Gaussian distribution from which mean ± S.D. were determined. (H) association and (I) dissociation kinetics rate constants of DNA cargo binding to DNA origami receptor with single binding sites measured by SPR. (J) Equilibrium constants, K*_D_*, for each site calculated as *k_d_*/*k_a_*.

To ensure that the DNA origami structure folded correctly, synthesis conditions were optimized, and DNA origami structures were visualized with transmission electron microscopy (TEM). DNA origami particles were clearly discernible in TEM micrographs and appeared to match the desired shape of the MSCE template with few (<5%) malformed structures observed (Figure 2E and Supplementary Figure 1). DNA origami structures tended to adsorb to the TEM grid surface along one of the particle’s four large flat sides. Given the design’s effective bilateral symmetry in two dimensions, it was impossible to distinguish between particles that landed on their right side versus those that landed on their left side. It was similarly impossible to distinguish between molecules that adsorbed on their top faces versus those that landed on their larger bottom face because the micrographs don’t provide height data. Thus, when 2D class averages were generated, the particles were classified in one of two orientations (Figure 2F). These allowed us to experimentally quantify the frequency distribution of each dimension of the MSCE template. Observed dimensions (mean ± S. D.) were within 10% of designed dimensions (Figure 2G) confirming that DNA origami MSCE templates formed the target structure with high fidelity and high yield.

We measured the binding kinetics for the interaction between the cargo molecule and each binding site independently with surface plasmon resonance (SPR). These experiments were performed using immobilized DNA origami receptor molecules containing one of either primary or secondary binding sites. Binding curves were consistent with a pseudo-first-order binding model (Supplementary Figure 2) from which binding rate constants and hence equilibrium dissociation constants could be reliably obtained (Figure 2H–J). Association rates (*k_a,1_* = 3.3 × 10^6^ ± 0.9 × 10^6^ M^-1^s^-1^ and *k_a,2_* = 1.6 × 10^6^± 0.8 × 10^6^ M^-1^s^-1^) and dissociation rates (*k_d,1_* = 4.0 × 10^-3^± 0.5 × 10^-3^s^-1^ and *k_d,2_* = 2.0 × 10^-2^± 0.1 × 10^-2^s^-1^) within the cavity of the DNA origami receptor were comparable but slower than previous measurements on a dextran-coated SPR chip^54^, which we attribute to differences in the local environment of DNA strands at the receptor binding site, each contributing different and ill-defined steric and electrostatic effects.

### Regular concentration-dependent exchange of DNA cargo

We first characterized a configuration of the DNA origami receptor with one primary and one secondary binding site (1:1). To quantify the expectations of the experiment it was useful to perform numerical simulations using an equivalent model previously presented by Åberg et al.^20^, except reformulated to explicitly contain the *C_eff_* term defined above. The remaining parameters in the numerical model are the kinetic rate constants of each of the binding sites: *k_a,1_* and *k_a,2_* for the association rate constants of primary and secondary binding sites respectively, and *k_d,1_* and *k_d,2_* for the dissociation rate constants of primary and secondary binding sites respectively (Supplementary Note 1). This reformulation thus allows all free parameters in the numerical model to be independently measured.

To demonstrate how *C_eff_* can be determined experimentally given independently measured binding rates, the effect of *C_eff_* on theoretical exchange rates (*k_exch_*) was determined from a first-order numerical model for a 1:1 system (Supplementary Note 1), parameterized with binding rates from SPR experiments are shown in Figure 3A. The shape of the plots and maximum exchange rates (*k_exch,max_*) are insensitive to *C_eff_*. In contrast, stability under dilution as quantified by the minimum exchange rate (*k_exch,min_*), and the range of [*C*] for which the system is sensitive to exchange, are both dependent on *C_eff_*. *C_eff_* is thus a key parameter for controlling the behavior of a 1:1 system. However, *C_eff_* is difficult to measure directly because it is affected by multiple entropic and enthalpic factors and hence cannot determined by the geometry of the system alone. However [*C*_50_], the concentration at which 50% change in exchange occurs (*k_exch_*([*C*_50_]) = 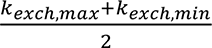), is a measurable parameter that increases linearly with *C_eff_* (Figure 3A inset). For the specific case where the kinetics of the primary and secondary binding sites are orthogonal but have the same binding kinetics (*k_a,1_* = *k_a,2_* = *k_a_*, *k_d,1_* = *k_d,2_* = *k_d_* and *K_D,1_* = *K_D,2_* = *K_D_*), this relationship is given by:

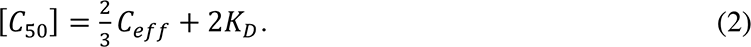

**Figure 3.**
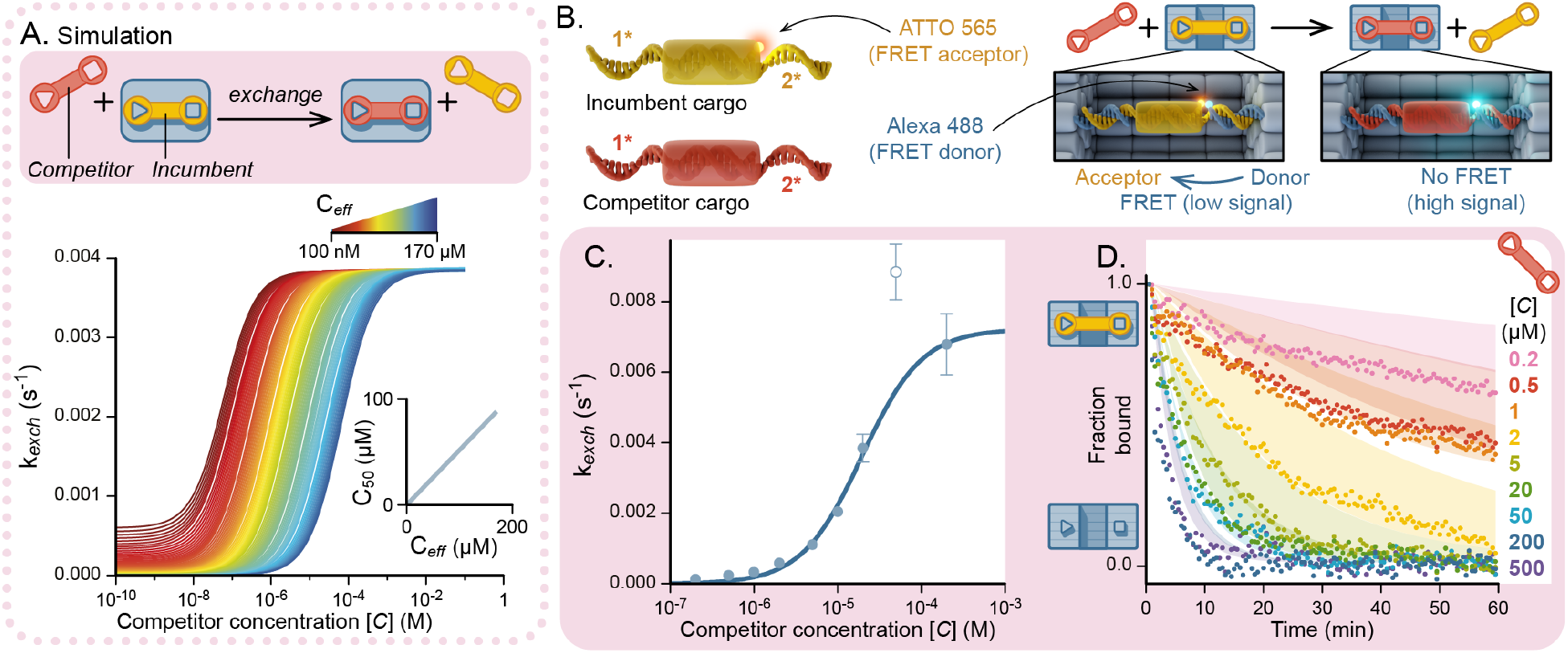
Concentration-dependent exchange of DNA cargo on a DNA origami receptor with a 1:1 configuration of binding sites. (A) Simulated dependence of exchange rate, *k_exch_*, of an incumbent cargo molecule on the concentration of competitor cargo molecules. *C_eff_* was varied from 0.1 *μM* to 170 *μM* (red to blue coloring). (B) The location of an acceptor fluorophore at the secondary binding site on the incumbent cargo molecule and a donor fluorophore at the secondary binding site on the receptor molecule for exchange measurements with FRET. Competitor cargo molecules were non-fluorescent. (C) Measured exchange rates from FRET measurements in the absence and presence of increasing concentrations of competitor molecule. Raw fluorescence intensity measurements are in Supplementary Figure 3. Error bars are mean +/- S.E.M for three independent repeats. One point (colored white) was removed from the sigmoidal fit by automated outlier detection (Q=1%) in GraphPad Prism. (D) Overlay of numerically predicted exchange kinetics and measured (dots) exchange kinetics. Numerical predictions are displayed as a shaded area corresponding to the area between minimum and maximum exchange rates that were within error of experimental measurements of binding site kinetics and *C_eff_*.

As described by Åberg *et al.*^20^ when *C_eff_* ≫ *K_D_*, which is likely for most feasible implementations then 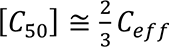. Similarly simple expressions can be used to calculate the maximum exchange rate 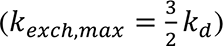 at high [*C*] and stability under dilution 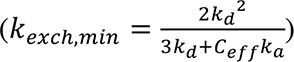, with equivalent primary and secondary binding sites ^15^. However, it is impractical to construct a real system with orthogonal binding sites that also have identical binding kinetics. We therefore derived general functional forms for *k*_*exch,min*_, *k_exch,max_* and [*C*_50_] in systems with one primary and one orthogonal secondary binding site, each with distinct kinetics (equations 13, 14 and 16 in Supplementary Note 2). These generalized functions similarly enable the *C_eff_* to be determined from [*C*_50_], given known rate constants of the primary and secondary site according to equation 3 below, which gives the solution for [*C*_50_] to leading order with the assumption *C_eff_* ≫ *K_D_*.

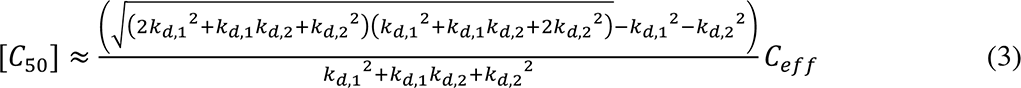

We next measured exchange experimentally on the 1:1 configuration of the DNA origami receptor with Förster resonance energy transfer (FRET). A donor fluorophore was located on the origami receptor near the secondary binding site (Figure 2B), and an acceptor fluorophore near the complementary site on pre-bound DNA cargo molecules (Figure 2C). Pre-bound receptor:cargo complexes were incubated at a concentration of 5 *nM* in the presence of 0.2 *μM* − 500 *μM* non-fluorescent DNA cargo. Consequently, exchange or dissociation could be measured as an increase in the intensity of donor fluorescence, which was monitored over time to obtain exchange kinetics. Measurements were also performed in the absence of competitor molecules and were used to normalize the donor fluorescence intensity with the assumption that these complexes do not dissociate at detectable levels on the timescales of the experiment. We observed concentration-dependent exchange, with normalized FRET data exhibiting an increase in exchange rate with increasing concentration of competitor cargo molecules (Figure 3B and Supplementary Figure 3). Since the concentration of competitor molecules [*C*] was large relative to the concentration of DNA origami receptor molecules in experiments where binding sites were in a 1:1 configuration, depletion of [*C*] and rebinding of incumbent cargo molecules was insignificant. Consequently, FRET dissociation curves in the 1:1 configuration were pseudo-first-order and fit with a single-exponential decay function (Supplementary Figure 3) to obtain exchange rates (*k_exch_*). When plotted against the competitor cargo concentration, *k_exch_* produced a sigmoidal shaped plot as predicted (Figure 3C). *k_exch_* vs [C] was then fit to a sigmoid function to heuristically determine *k_exch,max_*= 7.2 × 10^-3^ ± 0.4 × 10^-3^ s^-1^ and [*C*_50_] = 19.5 ± 0.1 µM. The minimum rate was set to 0 for the purposes of fitting as *k_exch,min_* was predicted to be below the detection limit in this experiment. Substituting into equation 3, this estimate of [*C*_50_] enables us to determine *C_eff_* to be 38 *μM*, which is similar to the concentration of one molecule in a volume of a sphere with radius equal to the length of the DNA cargo (∼50 *μM*). Along with SPR measurements, determining *C_eff_* fully parametrises the system of equations above, thus defining the equilibrium dissociation constant (*K_D_* = 0.4 ± 0.3 *pM*). In turn, the generalized equations for exchange rates in Supplementary Note 2 then allow the mean dissociation time under dilution of six days and a mean dissociation time at maximal exchange rate of four minutes to be calculated. Moreover, numerical models (Supplementary Note 1) could be solved with no free parameters, which yielded curves that were reasonably consistent with experimental data (Figure 3D).

### Multi-site concentration-dependent exchange of DNA cargo

Having demonstrated that the 1:1 configuration of the DNA origami receptor exhibits concentration-dependent exchange of DNA cargo, the system was expanded to a configuration consisting of a single secondary and two primary binding sites (2:1) (Figure 2A). This enables an additional exchange pathway via MSCE, resulting in a biphasic dependence of exchange rate on the competitor cargo concentration (Figure 4A, Supplementary Note 3) (Spinney *et al.* 2022). MSCE occurs in the first phase and when *K_D,1_* ≪ *C_eff_*, *k_exch_* saturates in an intermediate stable regime at a rate of

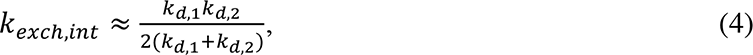

to leading order (Equation 25, Supplementary Note 2). The onset of the concentration-dependent increase in exchange rates varies depending on the individual site binding kinetics. We characterise this by defining the concentration, [*C*_50_], where *k_exch_* is halfway between the minimal rate and the saturating intermediate rate, such that *k_exch_*([*C*_50_]) = 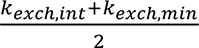. To leading order this is

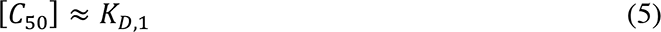

Thus, when *K_D,1_* ≪ *C_eff_*, [*C*_50_] is well approximated by *K*_D,1_ (Equation 27, Supplementary Note 2 and Supplementary Figure 4). Thus, competitive exchange can occur at competitor concentrations that are orders of magnitude lower than in the 1:1 configuration described above^20^. More rapid exchange occurs in the second phase when [*C*] exceeds *C_eff_*, and exchange is dominated by the kinetics of the 1:1 configuration.

**Figure 4.**
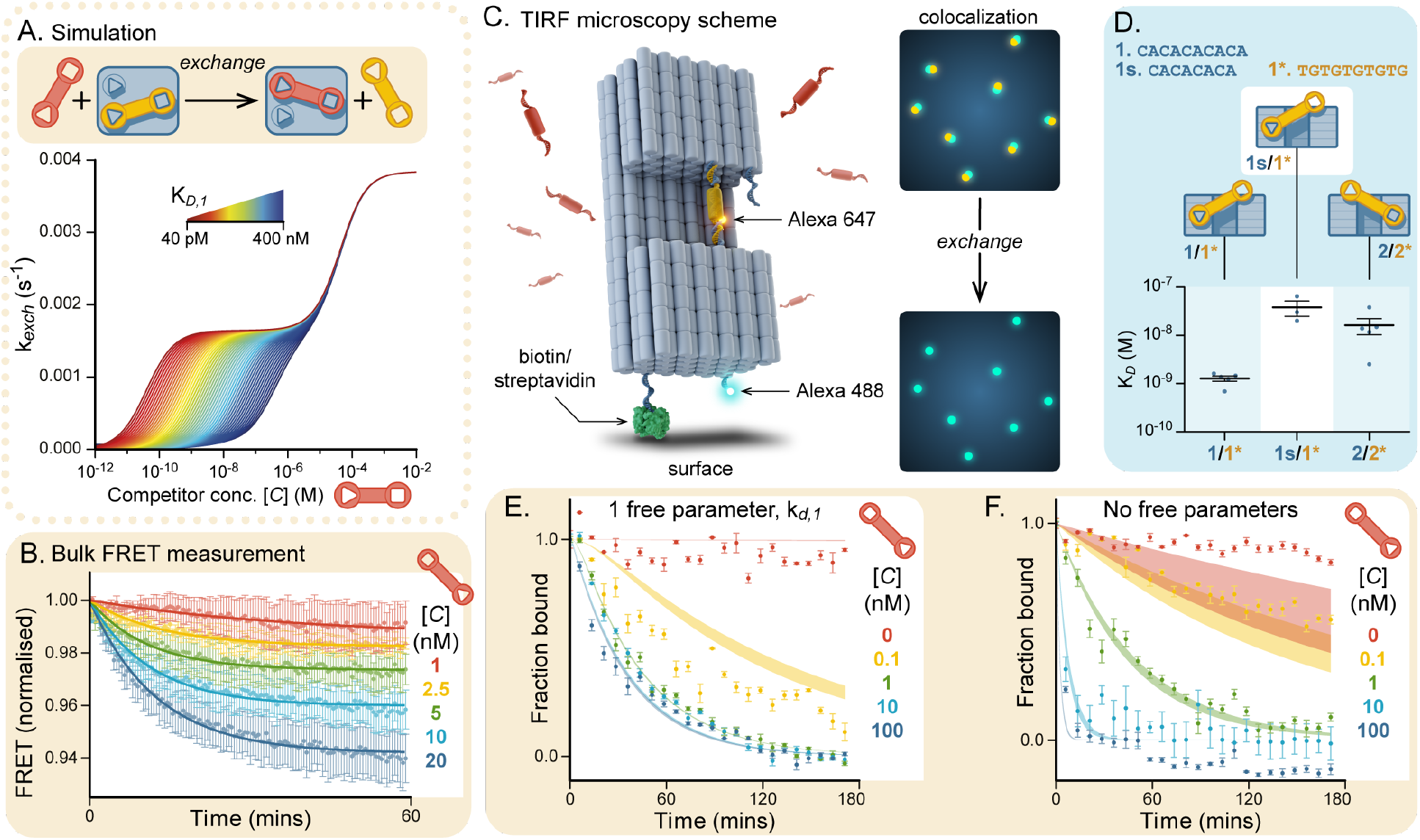
Multi-site concentration-dependent exchange of DNA cargo on a DNA origami receptor with a 2:1 configuration of binding sites. (A) Simulated dependence of exchange rate, *k_exch_*, of an incumbent cargo molecule, on the concentration of competitor cargo molecules. The equilibrium dissociation constant for the primary binding site, *K_D,1_*, was varied from 0.04 *nM* to 400 *nM* (red to blue coloring) while the effective concentration was kept constant, *C_eff_* = 38 *μM*. (B) FRET measurements of exchange of an incumbent Alexa-568 labeled DNA cargo molecule incubated in the absence and presence of increasing concentrations of competitor DNA cargo molecules. Error bars are mean ± S.E.M. for three independent repeats. (C) Conceptual depiction of TIRF microscopy experiment. (D) Equilibrium dissociation constant, K*_D_*, for the 8-nucleotide primary binding site consisting of 4 × *CA* repeats, 1s, as measured by SPR and compared to data for 10-nucleotide primary and secondary binding sites from Figure 2J. (E) Normalized kinetic data of subunit exchange from TIRF colocalization measurements in the absence and presence of increasing concentrations of competitor DNA cargo molecules containing two 10-nucleotide (CA)_5_ primary binding sites on the origami receptor fit to a numerical model with *k_d,1_* as a global free parameter. Bounds taken from mean ± S.D. of independently measured *k_a,1_*, *k_a,_*_2_, *k_d,_*_2_, *C_eff_*. RMSD=0.0071. (F) Normalized kinetic data for a DNA origami receptor containing two 8-nucleotide 4× *CA* primary binding sites fit to a numerical model with no free parameters and bounds again taken from mean ± S.D. of the independently measured rates and *C_eff_* (which was assumed to be the same as for the DNA origami receptor with 10-nucleotide primary binding sites). RMSD=0.0076. Error bars of experimental points are mean ± S.E.M. from three independent repeats.

Bulk FRET measurements demonstrate MSCE experimentally in the 2:1 configuration of the DNA origami receptor. As predicted, exchange occurred at concentrations orders of magnitude lower than required for exchange in the 1:1 configuration (Figure 4B). Subunit exchange measurements were therefore performed at lower concentrations of competitor molecules ([*C*] = 1 *nM* − 20 *nM*), similar to the concentration of DNA origami receptor molecules in FRET experiments ([*R*] = 5 *nM*). Exchange kinetics from FRET measurements in the 2:1 configuration were therefore second order, with significant depletion of competitor cargo concentrations and rebinding of incumbent fluorescent cargo molecules as the system approached equilibrium with different proportions of bound incumbent cargo molecules (Figure 4B). Second order analytical models for MSCE exchange kinetics are yet to be developed and likely to be substantially more complex than models describing first order exchange kinetics.

To directly relate experimental measurements of MSCE kinetics to existing first order models, we also measured exchange kinetics in a pseudo-first-order regime. This was achieved with surface-based single-molecule co-localization measurements using total internal reflection fluorescence (TIRF) microscopy. TIRF experiments enabled exchange measurements to be made at vanishingly low DNA origami concentrations, where depletion of DNA cargo concentrations and rebinding of incumbent cargo were insignificant, thus readily satisfying conditions for pseudo-first-order kinetics. For TIRF experiments, a biotinylated DNA strand was added to the base of the DNA origami receptor for immobilization on the surface of a BSA biotin:streptavidin-coated coverglass. Since competitor molecules were not fluorescently labelled, exchange was monitored as a reduction in colocalization of Alexa-488-labelled DNA origami receptor and Alexa-647-labeled incumbent cargo molecules as a function of time (Supplementary Figure 5). To reduce FRET the Alexa-488 fluorophore was moved away from the secondary binding site and to the base of the DNA origami receptor, increasing distance to the acceptor, and the Atto-565 fluorophore on the DNA cargo was replaced with an Alexa-647 fluorophore to reduce spectral overlap (Figure 4C).

Colocalization measurements of preformed receptor:cargo complexes were performed in the absence and presence of unlabeled DNA cargo at concentrations of 0, 0.1,1,10 and 100 nM over a period of three hours. The complex was stable under dilution with no evidence of spontaneous dissociation on the timescale of the experiment. In contrast, dissociation of the incumbent cargo was observed in the presence of all competitor cargo concentrations tested (Figure 4E). Surprisingly however, the saturating exchange rate was slower than the predicted rate of 0.002 s^-1^ (mean first passage time=10 mins). Indeed, a numerical model of MSCE kinetics (Supplementary Note 3) fully parameterized with experimentally measured binding kinetics and *C_eff_* was not consistent with measured exchange rates (Supplementary Figure 6).

We postulated that the single-stranded DNA extensions at each of the two primary binding sites in the 2:1 configuration may be sufficiently long to allow for cross-reactivity with a single DNA cargo molecule, which could significantly slow dissociation rates. We therefore performed a global fit of a numerical model of MSCE kinetics with the dissociation rate of the primary site, *k_d,1_*, as the single free parameter, which resulted in a good fit to the experimental data with a 4-fold decrease (bounds of fit 0.97-1.00×10^-3^ s^-1^) compared to SPR data (*mean* ± *S*. *D*. = 4.0 ± 0.5 × 10^-3^*s*^-1^). Mean first passage times can thus be calculated with equations in Supplementary Note 2, which gives a separation of timescales from 25 days to 35 minutes at the first plateau and a minimum mean exchange time of 17 minutes.

A crucial feature of this implementation is that particular elements of the design are addressable in order to readily elicit desired outcomes in terms of the exchange rate of the molecule of interest. To this end a DNA receptor was designed in which the primary binding site was shortened from 10 nucleotides to eight nucleotides by reducing the number of CA repeats to four. Reducing the stability of the primary binding site predicts a faster maximal exchange rate and an increase in the competitor concentration around which exchange is sensitive since from equation 5 above, *k_exch_*([*C*_50_]) ≈ *K_D,1_*. Moreover, by shortening the length of the binding strand, we sought to simultaneously minimize cross-reactivity between the two primary binding sites and thus allowing the kinetics of individual binding sites to be more consistent with independent measurements with SPR (Figure 4D and Supplementary Figure 7).

TIRF measurements revealed an increase in maximal exchange rate, which required a higher concentration of [*C*] as expected (Figure 4F). Moreover, the numerical model of MSCE kinetics, fully parameterized with experimental measurements was remarkably consistent with exchange rates, which in turn is consistent with cross reactivity confounding kinetic measurements in configurations with two 10-nucleotide primary binding sites. The dissociation under dilution was, however, slower than predicted, which may in part be due to the rebinding of dissociated incumbent cargo strands. Nonetheless, with the DNA origami receptor in the 2:1 configuration and 8-nucleotide primary binding sites, a large separation of timescales for mean exchange times was achieved, which ranged from several hours under dilution, to approximately one minute in the presence of competitor molecules. This demonstrates that with careful design and characterization, it is possible to build a fully predictable MSCE system, which we anticipate will be of general use in DNA nanotechnology.

### Multi-site concentration-dependent exchange of protein cargo

While toehold exchange has been used effectively as the primary mechanism for activating subunit exchange and actuating molecular motions in DNA nanostructures, toehold exchange requires DNA-based binding sites. MSCE in contrast is not limited to DNA-based cargo and hence can be used to facilitate the dynamic exchange of other biomolecules or nanoparticles. To demonstrate this generalizability, the DNA origami receptor was designed to be adaptable and to accommodate a generalized cargo molecule, and other chemical moieties can readily be incorporated to bind specifically to non-DNA cargo molecules. Here, a protein-based implementation is presented. Inspired by the exchange of DNA polymerase subunits in the replisome from which the MSCE model was developed, we used Klenow-fragment from *E. coli* DNA polymerase as the target cargo molecule for exchange. The protein was labelled with Alexa-647 fluorophore and expressed with 10× histidine peptides at both the N- and C-terminus, which bind specifically to 5× nickel-NTA motifs tethered to DNA strands^56^ at specific locations on the DNA origami receptor (Figure 5A). One nickel-NTA motif was located on one side of the cavity to act as a secondary binding site, and one or two nickel-NTA motifs were located on the other side of the cavity as primary binding sites. This arrangement allowed the protein to bind across the cavity of the DNA origami receptor, while being spaced too widely for the protein to readily bind across one side of the cavity. The receptor thus facilitates a 1:1 configuration for regular exchange (Figure 5B) of protein subunits and a 2:1 configuration that enables MSCE (Figure 5C).

**Figure 5:**
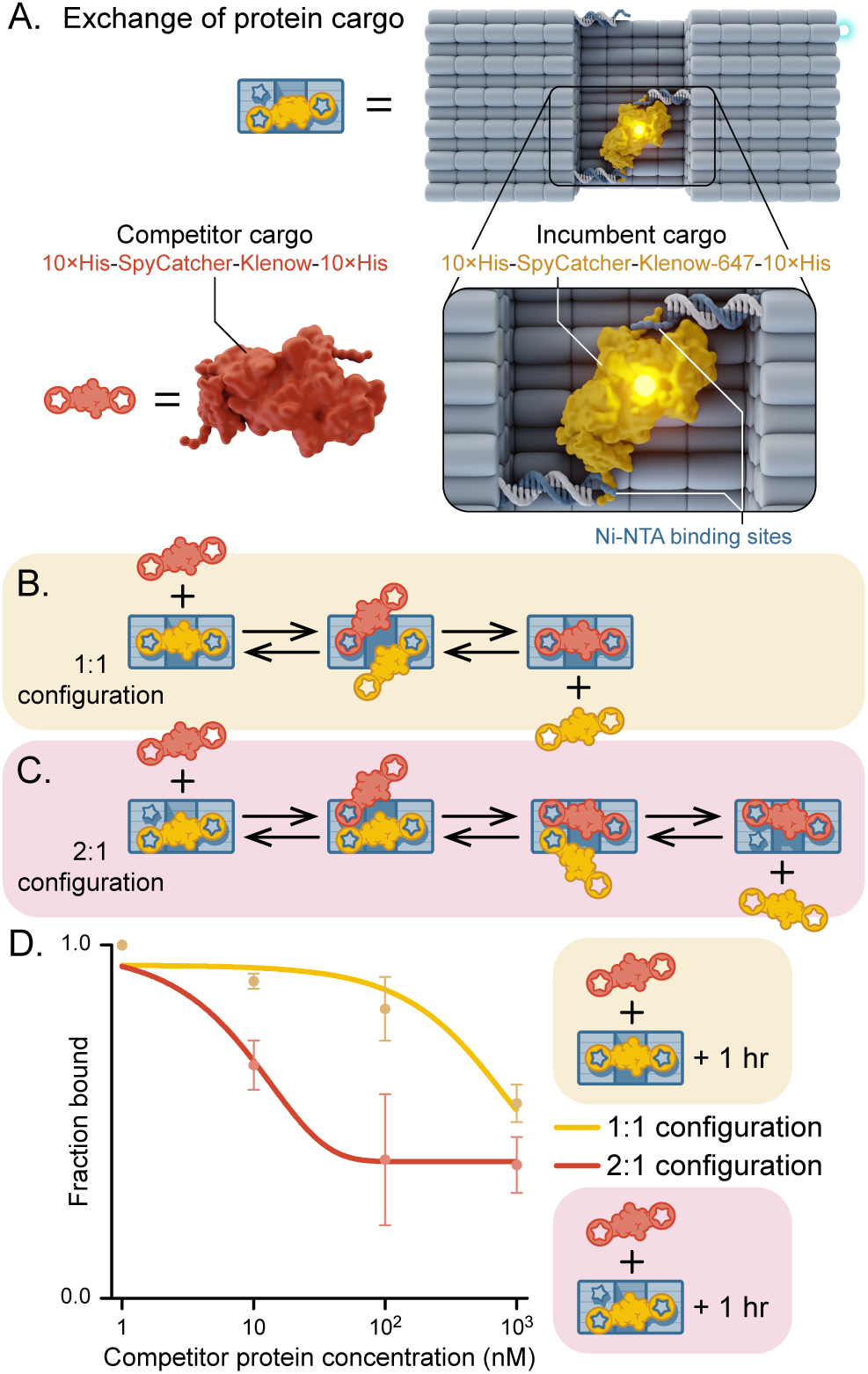
Multi-site concentration-dependent exchange of protein cargo. (A) Depiction of protein components added to DNA origami receptors to characterize protein-based exchange. (B) Protein exchange on a DNA origami receptor with a 1:1 configuration of binding sites. (C) Protein exchange on a DNA origami receptor with a 2:1 configuration of binding sites. (D) Normalized colocalization of Alexa488-labelled DNA origami receptor with Alexa647-labelled protein after incubation for one hour in the presence of unlabeled competitor protein at increasing concentrations. Error bars are mean ± S.E.M. for three independent repeats.

TIRF microscopy was again used to measure colocalization of Alexa-488 labeled DNA origami receptor molecules with Alexa-647 protein. To capture both regular exchange in the 1:1 configuration and MSCE in the 2:1 configuration, it was necessary to perform measurements over a large range of competitor concentrations (0, 10, 100 and 1000 *nM*). However, TIRF measurements in the presence of high competitor concentrations resulted in a detectable increase in background fluorescence. We attributed the high background to a substantial amount of non-specific binding of protein to the glass coverslip, which might confound kinetic measurements. Thus, Alexa-488 labeled receptor was co-purified with Alexa-647 labeled protein cargo, before being incubated in the presence of increasing concentrations of unlabeled protein cargo for 1 hour. Free protein was then rapidly removed by size exclusion spin-column chromatography before colocalization was measured with TIRF microscopy (Figure 5D). A concentration dependent decrease in the fraction of bound origami with fluorescently labeled protein cargo was observed in both the 1:1 and the 2:1 configurations of the DNA origami receptor. In the 2:1 configuration, exchange occurred at competitor concentrations that were orders of magnitude lower than in the 1:1 configuration illustrating MSCE of protein subunits.

This experiment thus demonstrates the more general application of the controlled exchange over time of non-DNA cargo molecules localized with nanometer precision on a DNA origami receptor, which we anticipate to be of broad applicability in the design of novel protein-based toehold exchange systems, and which greatly expands the available chemical space in these systems into the controllable localization and replacement of enzymes.

## CONCLUSIONS

We present the first engineered implementation of the MSCE mechanism, demonstrating the formation of stable complexes between a DNA origami receptor molecule and protein or DNA cargo molecules, which were capable of rapid exchange with competitor cargo molecules. The system exhibited predictable exchange kinetics that could be readily reconfigured or tuned to control stability and exchange rates. One configuration achieved stability-versus-exchange rates that were separated by over three orders of magnitude: spontaneous dissociation of cargo molecules under dilution required days, whereas exchange in the presence of nanomolar concentrations of competitor molecules occurred in minutes. Importantly, MSCE is not limited to DNA cargo and hence offers a stepwise advance over DNA toehold exchange. We demonstrate this generalizability of MSCE using a DNA polymerase protein as a cargo molecule to achieve rapid protein exchange within an otherwise stable complex. Finally, this study constitutes a new application for DNA nanostructures in the context of ‘designer kinetics’ via multivalent interactions, to engineer the interplay between stability and exchange in a rational way, potentially unlocking a wide range of potential synthetic soft systems at the nanoscale.

## METHODS

Numerical modelling are described in Supplementary Notes 1 and 3, and analytical models are described in Supplementary Note 2.

### DNA cargo design and synthesis

DNA cargo was designed in NUPACK^57^ with binding sites according to Hertel et.al.^54^ and consisted of two HPLC purified ssDNA strands (Integrated DNA Technologies (IDT). One strand, the binding strand, contained both binding sites 1* and 2* as well as a central region complementary to the second strand, termed the complementary strand. For fluorescently labelled DNA cargo molecules, one strand of the DNA duplex was purchased modified with Atto-565 or Alexa-647 as appropriate. These were annealed in a 1:1.5 ratio of binding to complementary strand in hybridization buffer (20 mM Tris HCl pH 8, 150 mM NaCl and 10 mM MgCl_2_) by rapidly heating the mixture to 95 °C in a thermal cycler for 2 min, and then cooling at a rate of −0.4 °C per 1 min to 25 °C. Synthesis was confirmed by native PAGE gel (Supplementary Figure 8).

Synthesized DNA cargo was stored at −20 °C for up to one year. DNA cargo was diluted in DNA imaging buffer (20mM Tris pH 8, 150 mM NaCl, 20 mM MgCl_2_, 3 mM EDTA, 0.005% Tween 20) for all experiments.

### Protein cargo design and synthesis

Protein cargo was designed as a SpyCatcher-Klenow(C907S) fusion protein separated by a glycine-serine-glycine (GSG) linker with both N-terminal and C-terminal 10×His-tags with a cysteine adjacent to each 10×His tag in a pET21b vector (Genscript). The plasmid was transformed into T7 express cells (NEB) according to manufacturer’s instructions and expressed in Luria-Bertani broth at 37 °C with shaking at 180 rpm before induction by IPTG at an optical density of 0.6, after which the temperature was lowered to 18 °C and the cultures were incubated overnight before centrifugation at 6000 rcf, 4 °C for 30 mins. Cell pellets were resuspended in buffer A (20 mM Tris-HCl (pH 8.0), 300 mM NaCl, 1 mM DTT, 1 tablet EDTA-free Protease Inhibitor, Roche) and lysed by probe sonicator (3× 3min, 50% cycles, 30% power, Branson Sonifier 250, Emerson) before centrifugation at 10,000 rcf, 4 °C for 20 mins. Supernatant was applied to a 5 ml HisTrap HP column (GE Healthcare) using an AKTA Prime liquid chromatography system before washing with buffer B (20 mM Tris-HCl (pH 8.0), 300 mM NaCl, 5 mM TCEP, 20 mM Imidazole) and elution with a gradient of buffer B and buffer C (20 mM Tris-HCl (pH 8.0), 300 mM NaCl, 5 mM TCEP, 1 M Imidazole). Samples of elution fractions were taken to perform sodium dodecyl sulfate polyacrylamide agarose gel electrophoresis (SDS-PAGE) (Bolt 4-12% Bis-Tris Plus, Invitrogen) at 180 V for 30 min in 1× MES buffer (Novex) using a SeeBlue Plus2 protein standard (Invitrogen) with Coomasie Brilliant Blue R-250 (Merk) post-staining. Fractions containing pure protein were collected and purified by size exclusion chromatography using a HiLoad 16/600 Superdex 200 pg column with elution in buffer D (20 mM Tris-HCl (pH 8.0), 300 mM NaCl, 5 mM TCEP). Samples of elution fractions were taken to perform SDS-PAGE as above, fractions containing pure protein were pooled and the final concentration was determined by Direct Detect infrared spectrometry (Merk). Labelling with Alexa-647 maleimide (ThermoFisher) was performed according to the manufacturer’s instructions and size exclusion chromatography performed as above in buffer E (20 mM Tris-HCl (pH 8.0), 300 mM NaCl, 12.5 mM MgCl_2_, 0.005% Tween 20). SDS-PAGE was performed to determine labelling efficiency and labelled protein was stored at −80 °C. For non-fluorescent protein, N-ethylmaleimide was used in place of Alexa-647 and protein purification was performed as above.

### DNA receptor design and synthesis

5× Ni-NTA labelled DNA strands were prepared and PAGE purified from DNA oligos with 5 C_6_-amine modifications (IDT) as described previously^56^.

The DNA origami receptor was designed in caDNAno^58^ and 3D models produced using CanDo^59^ and Blender (www.blender.org). The caDNAno file detailing the design of the DNA origami receptor along with all DNA staple sequences in a Microsoft Excel spreadsheet are in associated online content. 10-100 nM M13mp18 scaffold (Bayou Biolabs) was incubated with an excess of staple strands (IDT). All staple strands were desalting purified and incubated in 5× excess of scaffold, except for the appropriate (primary, secondary or 5× Ni/NTA) binding sites (10× excess), and if used the biotin and/or Alexa-488 conjugated strands (25× excess) which were HPLC purified. If used, the pre-assembled DNA cargo was incubated at 150× excess.

Incubation was performed in origami synthesis buffer (5 mM Tris base, 1 mM EDTA, 5 mM NaCl and 30 mM MgCl_2_) in a thermal cycler with rapid heating to 65 °C for 15 min, before cooling at a rate of −0.1 °C per 1 min to 60 °C, and then −0.1 °C per 10 min to 40 °C, then −0.1 °C per 1 min to 25 °C. The reaction mixtures were then mixed 1:1 (v/v) with PEG precipitation buffer (15% PEG 8000 (w/v), 5 mM Tris, 1 mM EDTA, and 505 mM NaCl) before centrifugation at 14,500 rpm at room temperature for 30 min. The supernatant was removed immediately, and the remaining pellet was resuspended in origami synthesis buffer. DNA origami template was diluted in DNA imaging buffer for DNA cargo experiments and buffer E for protein cargo experiments.

For TIRFM experiments the synthesis mix included staples that allowed for labelling of the structure with an Alexa488 fluorophore and a biotin molecule, both at the base of the structure. For SPR experiments, neither of these staple strands were included in the synthesis mix and the staples used for the biotinylation of the DNA origami template were instead used to anchor the structure on the SPR chip. For FRET experiments, none of these staples were used and a new fluorescently labelled staple was included inside the cavity of the DNA origami template (Figure 1).

### Transmission electron microscopy

Purified DNA origami templates were absorbed for 5 min onto Formvar carbon coated copper grids which had been glow discharged for 60 s. The sample was wicked away using filter paper, the grids stained with 2% uranyl acetate, wicked immediately, then left to dry at room temperature. Micrographs were acquired using bright field on a FEI Tecnai G2 20 TEM with a LaB6 filament and BM Eagle digital camera. Measurements of DNA origami template were obtained manually using the line tool in Fiji^60^ and 2D class averaging was performed using Relion^55^.

### Surface plasmon resonance acquisition and analysis

Experiments were performed on a Biacore S200 instrument (GE Healthcare Life Sciences). All experiments were performed at room temperature using DNA imaging buffer and a flow rate of 10 µl/min.

CM3 sensor chips were coupled with streptavidin to near saturation (typically between 4000 and 7000 RU) using the amine coupling kit (GE Healthcare Life Sciences). After streptavidin was coupled, biotinylated DNA strands were injected into reference and experimental flow cells before excess biotin-binding sites were blocked with biotin in DNA imaging buffer. Experimental flow cells were then loaded with DNA origami template containing either binding site 1, 1s or 2. Experiments were performed with one of each binding sites across three flow cells on a chip, with the remaining flow cell being a reference cell with no DNA origami template. The surface was then conditioned with 2 injections of DNA imaging buffer followed by the indicated concentrations of DNA cargo. Reference-subtracted data are shown, and all data were fit to a single-exponential in MATLAB 2020b.

### FRET Data Acquisition and Analysis

FRET experiments were performed in a 384-well, square bottom, low volume plate with a sample volume of 25 μl using a FLUOstar Omega microplate reader (BMG Labtech) using the bottom optic with a 485/12 nm bandpass excitation filter and a 520/30 nm bandpass emission filter. 5 nM of purified Alexa-488 labelled DNA origami template pre-assembled with Atto-565 labelled DNA cargo in DNA imaging buffer were incubated with 0-200 μM non-fluorescent competitor DNA cargo. Data were collected at 30 s intervals, with 200 flashes per well per cycle and double orbital shaking for 10 s at 500 rpm between timepoints to prevent sample settling in the well. Five concentrations of competitor DNA molecules and a buffer only control were measured for each experiment and three or more replicates were performed for each concentration.

Fluorescence time traces acquired in the absence of competitor DNA cargo were subtracted from all traces in the presence of competitor cargo, and traces were normalized with t=0 set as 1. Initially direct numerical integration was performed to acquire *k_exch_*, however variations in the baseline due to systematic instrument noise led to large errors for higher rates, and under-sampling on the timescale of this experiment led to an overestimate of *k_exch_* at lower rates (Supplementary Figure 3C). For Figure 2E and Supplementary Figure 4 data were fit to a single exponential and the reciprocal time constant reported as *k_exch_*. For Figure 2F data the baseline of each time trace was set to 0 if a baseline was reached and otherwise was subtracted from the mean value of the baselines in that experiment. Parameter-free fitting was performed using custom code utilizing the ODE15s solver for the set of differential equations in supplementary note 1 in MATLAB 2020b using parameters defined in the main text with mean ± S. D. as upper and lower bounds for each parameter.

### TIRF Microscopy Data Acquisition and Analysis

Microfluidic devices and coverslips were prepared as described previously^61^. Images were collected on a custom built TIRF microscope described in Bereja et. al.^61^ with a power density of ∼1–3 W cm^-2^ (measured at the objective with the laser beam normal to the surface of the coverslip. Single particle photobleaching was first performed using purified, pre-assembled DNA origami template and DNA cargo in DNA imaging buffer immobilized on a coverslip to a density of ∼1000 particles per field of view (FOV). Images were acquired with alternating 488 nm and 647 nm excitation (20 mW) with a 100 ms exposure time and 300-800 frames per FOV. 3 independent repeats of 5 FOV were obtained. Images were analysed with the JIM-Immobilized-Microscopy-Suite (https://github.com/lilbutsa/JIM-Immobilized-Microscopy-Suite) and thresholds for single-particles obtained by fitting to a Gaussian distribution. Thresholds were set from single-particle detection as mean ± 2 s. d. of the photobleaching step height and used for detection of single particles in exchange experiments.

DNA cargo exchange experiments were then performed in which pre-assembled DNA origami template and DNA cargo were adhered to the surface before washing with DNA imaging buffer and incubation with competitor cargo at varying concentrations as described in the main text. Exchange experiments were performed under identical imaging conditions to single particle photobleaching except each FOV was acquired for 10 frames. 24 time-points were acquired at 7.5 min intervals and different FOVs were obtained for each time-point to minimize the effect of photobleaching.

Images were analyzed using the JIM-Immobilised-Microscopy-Suite to detect single particles in each of the 488 nm and 647 nm channels and to extract fluorescence intensity traces. The mean of these traces for each particle was calculated and colocalization analysis was performed using the thresholds determined by single-particle photobleaching. Colocalization was calculated as

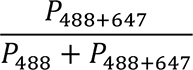

Where *P*_488_ is the number of particles within the single particle thresholds in the 488 channel but below the mean − 2 s. d. single particle threshold in the 647 channel and *P*_488+647_ is the number of particles within the single particle thresholds for both channels. This calculates the fraction of 488-labelled DNA-origami which is bound to 647-labelled incumbent cargo. This is shown graphically in supplementary figure 5. Chance coincidence was calculated as 3±1% by rotating the 647 channel by 90° and performing analysis as above.

Colocalization data for each competitor concentration were then normalized with the fraction bound at t=0 set to 1 and the fraction bound once a baseline was reached set to 0. This baseline was around 15% as shown in Supplementary Figure 5, which we attributed to a combination of chance coincidence and non-exchange competent DNA origami molecules which could be due to a proportion of molecules that had not fully integrated both primary binding sites. Competitor concentrations for which a baseline was not reached were normalized to the mean baseline of competitor concentrations for which a baseline was reached in that experiment.

Parameter-free fitting was performed using custom code utilizing the ODE15s solver for the set of differential equations in supplementary note 1 in MATLAB 2020b using parameters defined in the main text with mean ± S. D. as upper and lower bounds for each parameter. For parameter fitting, an optimisation problem was set up and solved using the nonlinear least-squares solver in MATLAB 2020b with k*_d,1_* as a single global free parameter and all other parameters as defined in the main text with mean ± S. D. as upper and lower bounds for each parameter.

Protein cargo exchange experiments were performed by incubating 100 nM DNA origami with 300nM Alexa-647 labelled protein cargo for 1h before addition to washed s300 resin (according to manufacturer’s instructions, GE Life Sciences) in a spin column and centrifugation for 4 mins at 1000 g to remove free protein. DNA origami:protein complex concentration in the flow through was measured by absorbance at 260 nm and diluted to 5 nM in buffer E before titration with the indicated concentrations of competitor protein for 1 h. Samples were then added to washed s300 resin in a spin column and centrifuged for 2 mins at 1000g, flow through was collected and the s300 purification was repeated to ensure removal of free protein. Samples were diluted to ∼100 pM for addition to a microfluidic device on a coverslip and immediate imaging as described above, except that 20 FOVs were obtained for one single timepoint. Colocalization analysis was performed as described above and normalized to the highest colocalization value in each experiment.

## ASSOCIATED CONTENT

Supporting information is available free of charge *via* the Internet at http://pubs.acs.org. These are incorporated in two documents (PDF). The first document consists of supplementary figures. The second document consists of supplementary notes detailing numerical models and analytical derivations. We also include details of the DNA origami design in an associated caDNAno file (JSON) and all DNA staple sequences in an associated spreadsheet (XLS).

## AUTHOR INFORMATION

### Corresponding Author

*Lawrence K. Lee, EMBL Australia Node for Single Molecule Science, School of Medical Sciences, UNSW Sydney, 2052, Australia.

### Author Contributions

The manuscript was written through contributions of all authors. All authors have given approval to the final version of the manuscript. ^^^These authors contributed equally.

## ACKNOWLEDGMENTS

The authors thank the facilities of Microscopy Australia at the Electron Microscope Unit, Mark Wainwright Analytical Centre, UNSW Sydney.

