## Supplementary for "Rapid exchange of stably bound protein and DNA cargo on a DNA origami receptor"

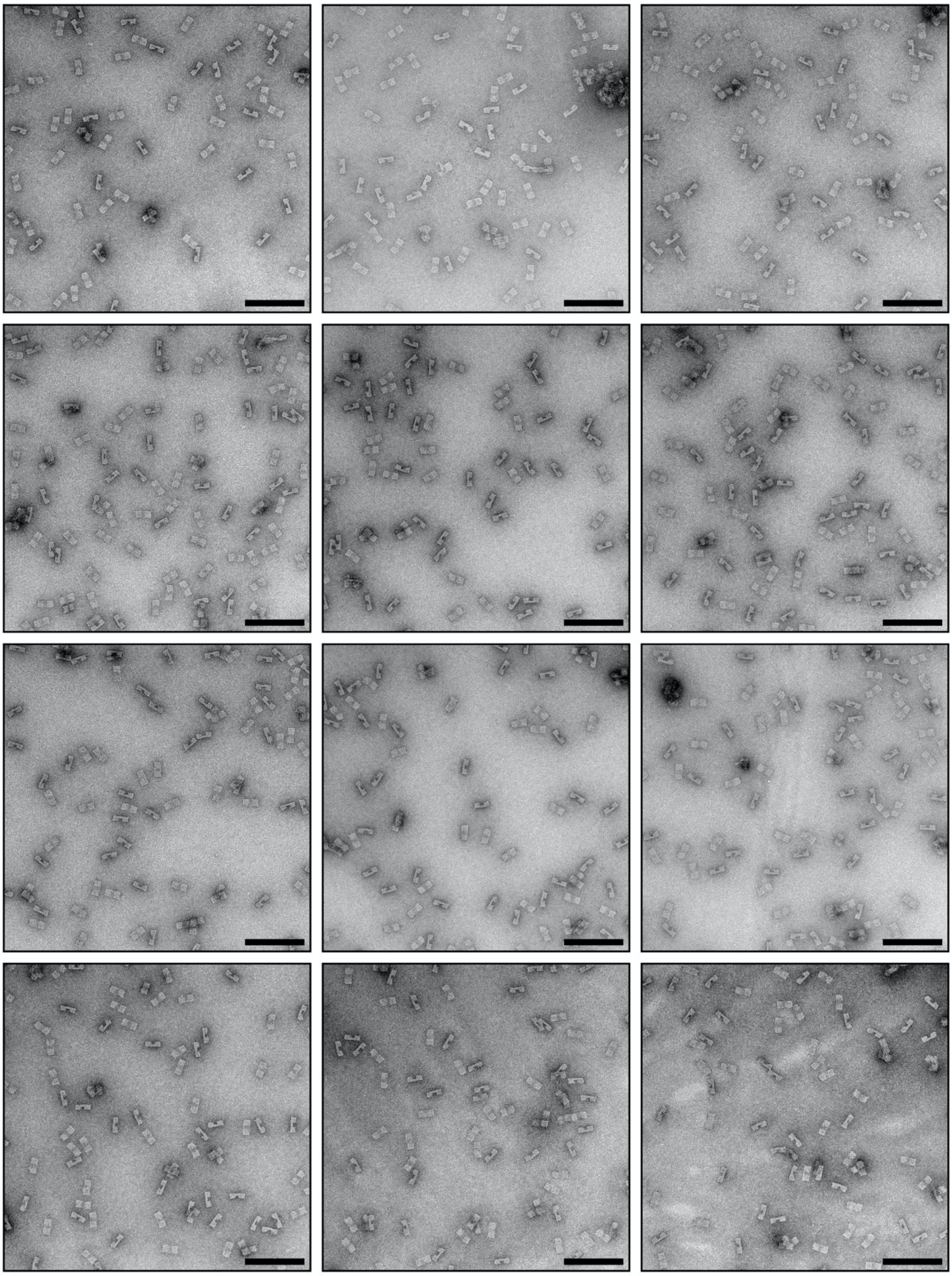

**Supplementary Figure 1. Exemplary TEM micrographs of DNA origami receptor molecules.**
Scale bar is 200 nm

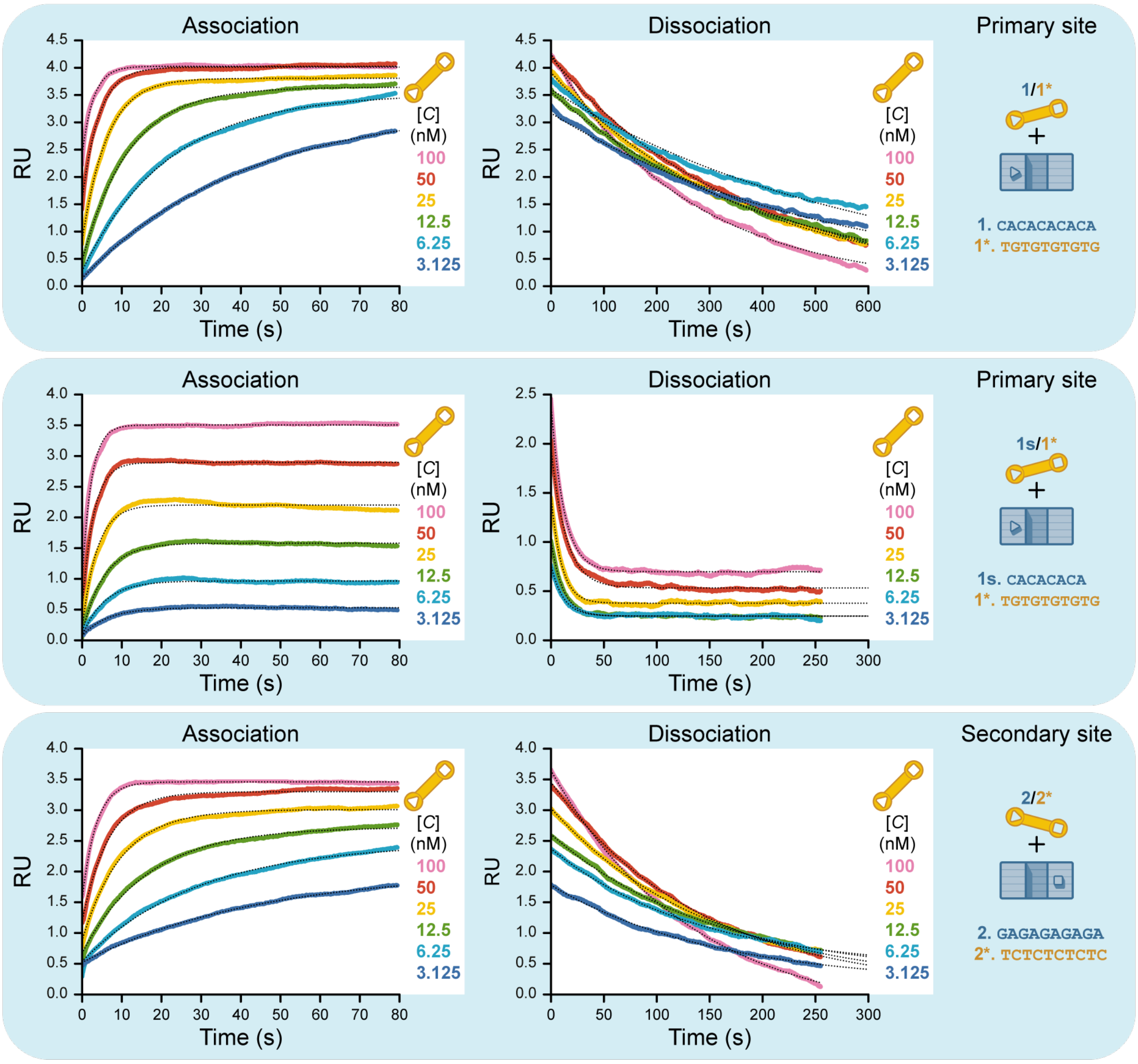


**Supplementary Figure 2. Exemplary surface plasmon resonance data.** Raw response units (RU), which are proportional to bound material, are shown for association (left) and dissociation (right) as dotted black lines. Association curves were fit separately for each concentration with a monoexponential equation (colored lines) to obtain an observed association rate, $k_{obs}$, according to: $RU_t = (RU_{max} - RU_0)(1 - e^{-k_{obs}t}) + RU_0$, where $RU_{max}$ and $RU_0$ are the maximal and initial response units and $t$ is time in seconds. Association rate constants, $k_a$, were then determined from the gradient of a linear function fit to plots of $k_{obs}$ vs $[C]$. Dissociation curves were fit to a monoexponential (colored lines) equation to obtain dissociation rate constants, $k_d$ according to: $RU = RU_0 e^{-k_d t} + RU_{min}$, where $RU_{min}$ are the minimum response units.

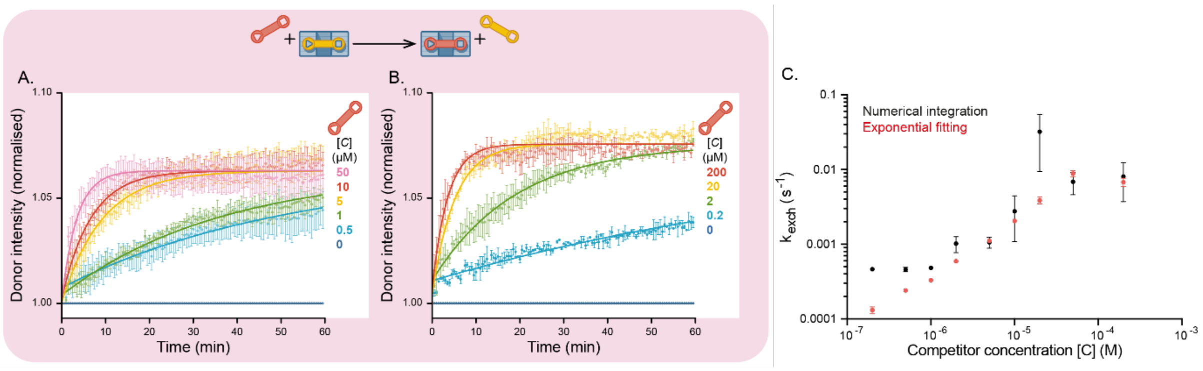


**Supplementary Figure 3. FRET data for cargo exchange on a DNA origami receptor with a 1:1 configuration of binding sites.** (A,B) FRET-based measurements of normalized Alexa-488 intensity in the absence and presence of increasing concentrations of competitor molecule. An increase in donor intensity is inversely proportional to a reduction in bulk FRET efficiency over time. Error bars are mean +/- SEM for 3 independent repeats. FRET data are fit with a monoexponential equation (solid lines) to estimate exchange rates, $k_{exch}$, according to $I = -I_{max}e^{k_{exch}t} + I_0$ where $I$ are the FRET intensity units, $I_{max}$ and $I_0$ are the maximum predicted and the initial FRET intensity units respectively. It was also possible to determine $k_{exch}$ by numerical integration of data, with a comparison of rates determined from numerical integration of data to exponential fitting shown in (C) (see Materials and Methods).

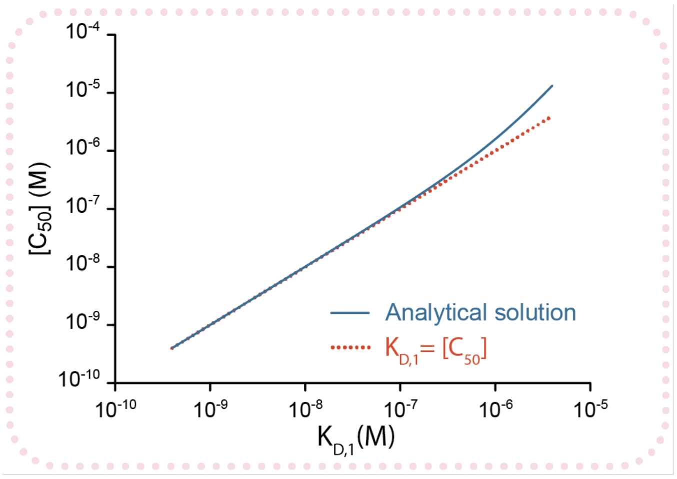


**Supplementary Figure 4. $[C_{50}]$ is well approximated by $K_{D,1}$ for a receptor with a 2:1 binding configuration.** A plot of $[C_{50}]$ vs $K_{D,1}$ according to analytical derivation (Equation 27, Supplementary Note 2) in blue compared with a line of equality in orange illustrates how $[C_{50}]$ is well approximated by $K_{D,1}$ when $K_{D,1} \ll C_{eff}$.

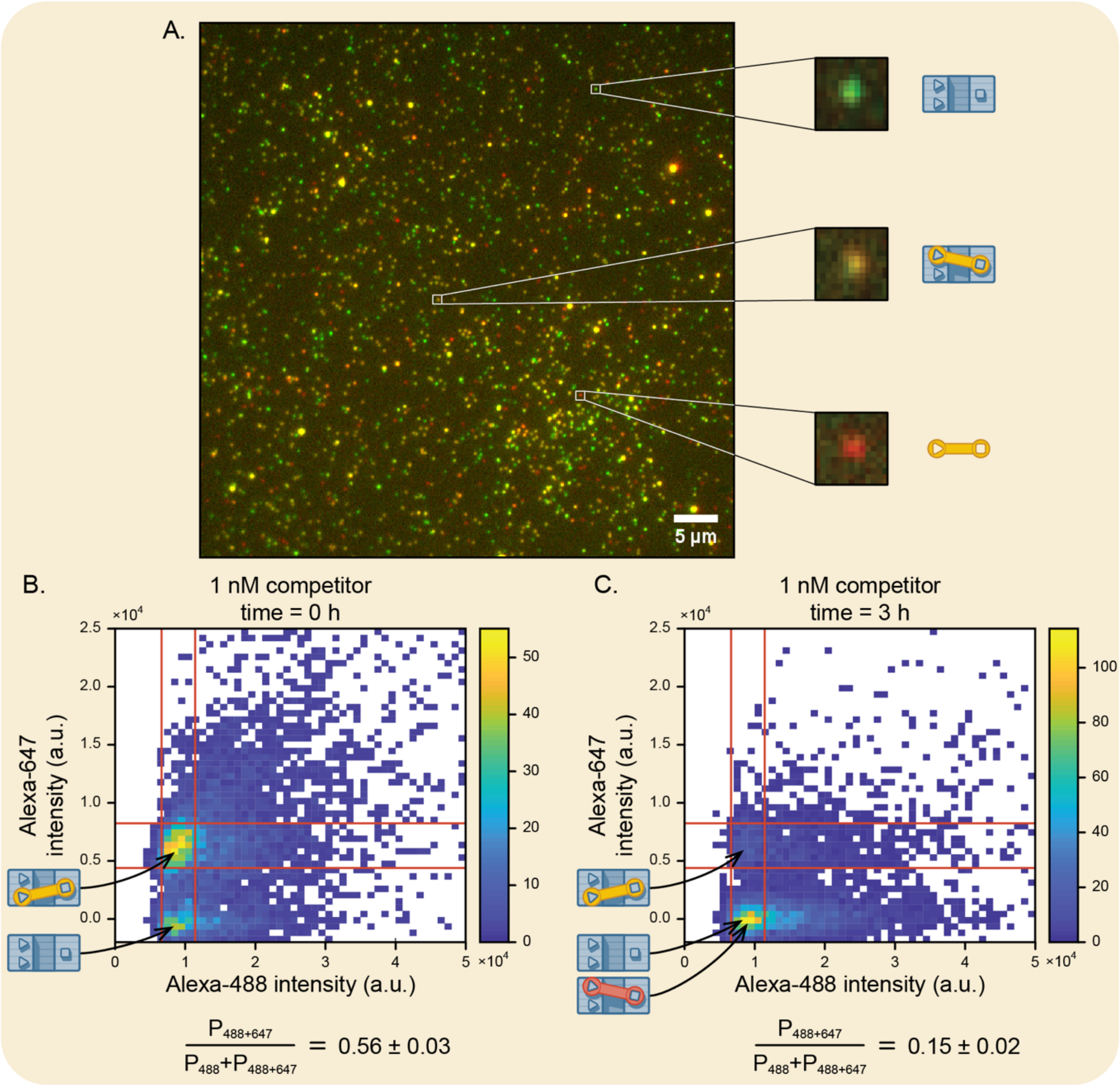


**Supplementary Figure 5. Example data from total internal fluorescence microscopy (TIRFM) measurements to illustrate data processing.** TIRFM was used to measure colocalization of DNA origami receptor (green) with pre-bound DNA cargo (red) in the presence of non-fluorescent competitor DNA cargo. (A) Representative TIRFM image of DNA cargo bound to DNA origami receptor, with example spots showing DNA receptor alone, co-localized with fluorescent cargo molecule, and a single isolated fluorescent cargo molecule. (B) 2D histogram of individual bound molecules (between red cutoffs determined as described in Methods) at time=0 in the presence of 1 nM competitor DNA cargo. (C) 2D histogram of individual bound molecules at time=3h in the presence of 1 nM competitor DNA cargo.

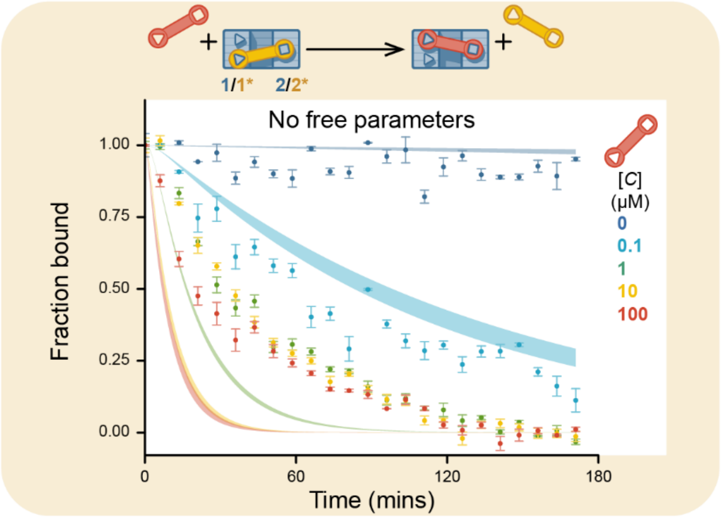


**Supplementary Figure 6. Normalized kinetic exchange data from TIRFM colocalization measurements on DNA origami receptors in a 2:1 configuration with 10-base $(GA)_5$ primary binding sites.** Data are shown fitted to a numerical model (Supplementary Note 3) with no free parameters. Bounds taken from mean ± S.D. of independently measured $C_{eff}$, and association and dissociation rate constants for each binding site. RMSD=0.0147.

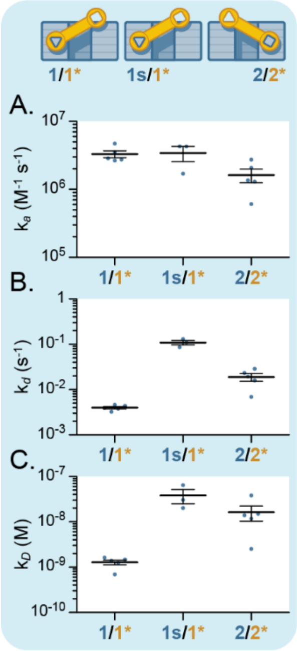


**Supplementary Figure 7. Kinetic measurements of different binding sites.** (A) Association and (B) dissociation kinetics rate constants of DNA cargo binding to DNA origami receptor at different binding sites as measured by SPR. (C) Equilibrium constants $K_D$ for each site, calculated as $k_d/k_a$.

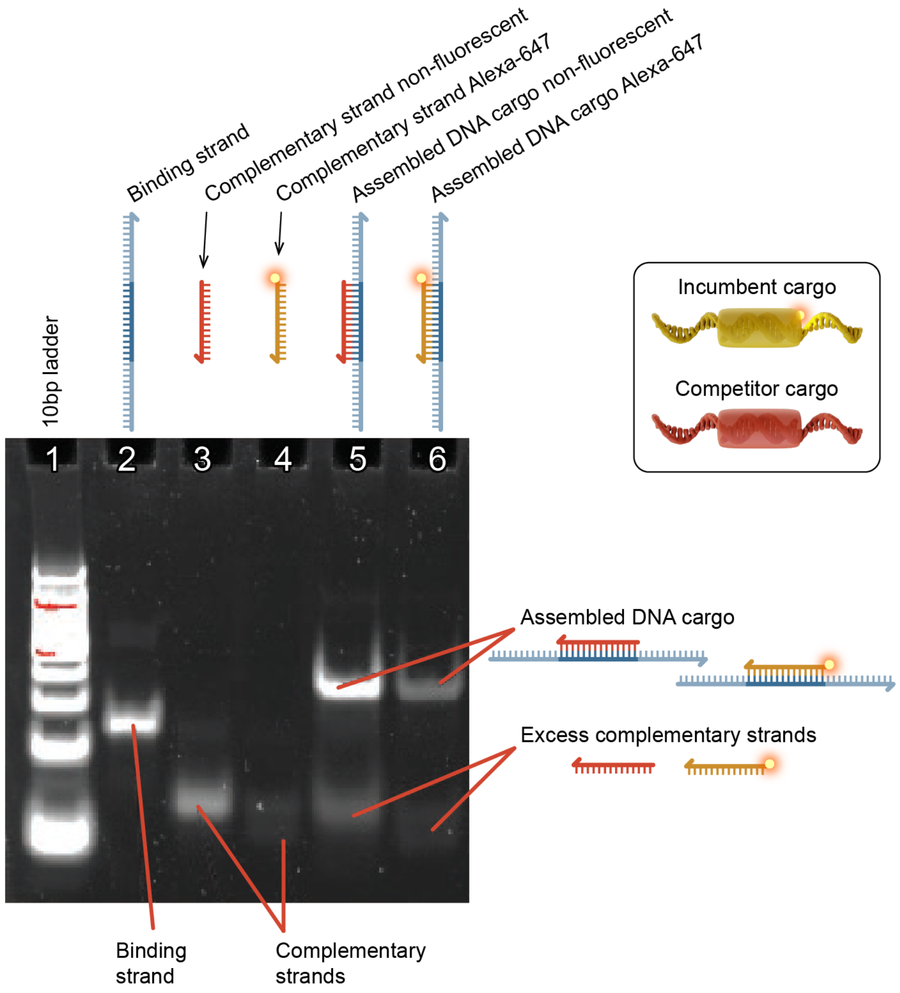


**Supplementary Figure 8. Native PAGE gel of synthesized binding and complementary fluorescent and non-fluorescent ssDNA strands.** The binding and complementary strands were in a 1:1.5 ratio during synthesis of the DNA cargo.
